# MutantHuntWGS: A Pipeline for Identifying *Saccharomyces cerevisiae* Mutations

**DOI:** 10.1101/2020.05.16.099259

**Authors:** Mitchell A. Ellison, Jennifer L. Walker, Patrick J. Ropp, Jacob D. Durrant, Karen M. Arndt

## Abstract

MutantHuntWGS is a user-friendly pipeline for analyzing *Saccharomyces cerevisiae* whole-genome sequencing data. It uses available open-source programs to: (1) perform sequence alignments for paired and single-end reads, (2) call variants, and (3) predict variant effect and severity. MutantHuntWGS outputs a shortlist of variants while also enabling access to all intermediate files. To demonstrate its utility, we use MutantHuntWGS to assess multiple published datasets; in all cases, it detects the same causal variants reported in the literature. To encourage broad adoption and promote reproducibility, we distribute a containerized version of the MutantHuntWGS pipeline that allows users to install and analyze data with only two commands. The MutantHuntWGS software and documentation can be downloaded free of charge from https://github.com/mae92/MutantHuntWGS.

## INTRODUCTION

*Saccharomyces cerevisiae* is a powerful model system for understanding the complex processes that direct cellular function and underpin many human diseases (Birkeland *et al*. 2010; Botstein and Fink 2011; Kachroo *et al*. 2015; Hamza *et al*. 2015, 2020; Wangler *et al*. 2017; Strynatka *et al*. 2018). Mutant hunts (*i*.*e*. genetic screens and selections) in yeast have played a vital role in the discovery of many gene functions and interactions (Winston and Koshland 2016). A classical mutant hunt produces a phenotypically distinct colony derived from an individual yeast cell with at most a small number of causative mutations. However, identifying these mutations using traditional genetic methods (Lundblad 1989) can be difficult and time-consuming (Gopalakrishnan and Winston 2019).

Whole-genome sequencing (WGS) is a powerful tool for rapidly identifying mutations that underlie mutant phenotypes (Smith and Quinlan 2008; Irvine *et al*. 2009; Birkeland *et al*. 2010). As sequencing technologies improve, the method is becoming more popular and cost-effective (Shendure and Ji 2008; Mardis 2013). WGS is particularly powerful when used in conjunction with lab-evolution (Goldgof *et al*. 2016; Ottilie *et al*. 2017) or mutant-hunt experiments, both with (Birkeland *et al*. 2010; Reavey *et al*. 2015) and without (Gopalakrishnan *et al*. 2019) bulk segregant analysis.

Analysis methods that identify sequence variants from WGS data can be complicated and often require bioinformatics expertise, limiting the number of investigators who can pursue these experiments. There is a need for an easy-to-use, data-transparent tool that allows users with limited bioinformatics training to identify sequence variants relative to a reference genome. To address this need, we created MutantHuntWGS, a bioinformatics pipeline that processes data from WGS experiments conducted in *S. cerevisiae*. MutantHuntWGS first identifies sequence variants in both control and experimental (*i*.*e*. mutant) samples, relative to a reference genome. Next, it filters out variants that are found in both the control and experimental samples while applying a variant quality score-cutoff. Finally, the remaining variants are annotated with information such as the affected gene and the predicted impact on gene expression and function. The program also allows the user to inspect all relevant intermediate and output files.

To enable quick and easy installation and to ensure reproducibility, we incorporated MutantHuntWGS into a Docker container (https://hub.docker.com/repository/docker/mellison/mutant_hunt_wgs). With a single command, users can download and install the software. A second command runs the analysis, performing all steps described above. MutantHuntWGS allows researchers to leverage WGS for the efficient identification of causal mutations, regardless of bioinformatics experience.

## METHODS

### Pipeline Overview

The MutantHuntWGS pipeline integrates a series of open-source bioinformatics tools and Unix commands that accept raw sequencing reads (compressed FASTQ format or .fastq.gz) and a text file containing ploidy information as input, and produces a list of sequence variants as output. The user must provide input data from at least two strains: a control strain and one or more experimental strains. The pipeline uses (1) Bowtie2 to align the reads in each input sample to the reference genome (Langmead and Salzberg 2012), (2) SAMtools to process the data and calculate genotype likelihoods (Li *et al*. 2009), (3) BCFtools to call variants (Li *et al*. 2009), (4) VCFtools (Danecek *et al*. 2011) and custom shell commands to compare variants found in experimental and control strains, and (5) SnpEff (Cingolani *et al*. 2012) and SIFT (Vaser *et al*. 2016) to assess where variants are found in relation to annotated genes and the potential impact on the expression and function of the affected gene products (Figure 1). A detailed description of the commands used in the pipeline and all code is available on the MutantHuntWGS Git repository (https://github.com/mae92/MutantHuntWGS; see README.md, Supplemental_Methods.docx files).

**Figure 1.**
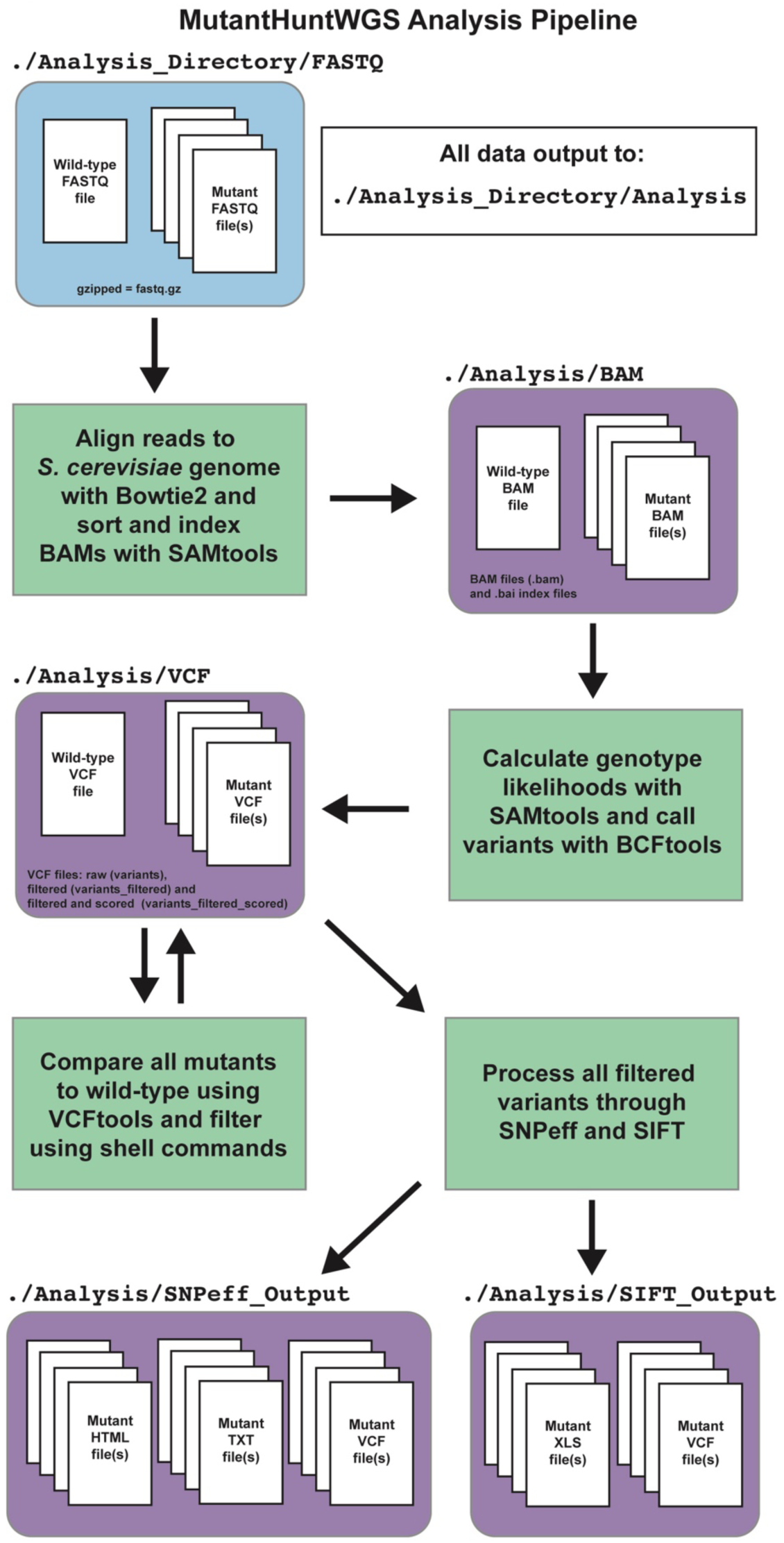
Flow chart of the MutantHuntWGS pipeline. Input data are colored in blue, the various bioinformatics tools in the pipeline are colored in green, and output data are colored in purple. Arrows identify the path of the workflow at each step of the pipeline.

### Analysis of previously published data

To demonstrate utility, we used MutantHuntWGS to analyze published datasets from paired-end sequencing experiments with DNA prepared from bulk segregants or lab-evolved strains (Birkeland *et al*. 2010; Goldgof *et al*. 2016; Ottilie *et al*. 2017). These data were downloaded from the sequence read archive (SRA) database (https://www.ncbi.nlm.nih.gov/sra; project accessions: SRP003355, SRP074482, SRP074623) and decompressed using the SRA toolkit (https://github.com/ncbi/sra-tools/wiki). MutantHuntWGS was run from within the Docker container, and each published mutant (experimental) file was compared to its respective published control.

### Data Availability Statement

All code and supplementary information on the methods used herein are available on the MutantHuntWGS Git repository (https://github.com/mae92/MutantHuntWGS).

## RESULTS

### Installing and running the MutantHuntWGS pipeline

To facilitate distribution and maximize reproducibility, we implemented MutantHuntWGS in a Docker container (Boettiger 2015; Di Tommaso *et al*. 2015). The container houses the pipeline and all of its dependencies in a Unix/Linux environment. To download and install the MutantHuntWGS Docker container, users need only install Docker Desktop (https://docs.docker.com/get-docker/), open a command-line terminal, and execute the following command:

~~~
$ docker run -it -v
/PATH_TO_DESKTOP/Analysis_Directory:/Main/Analysis_Directory mellison/mutant_hunt_wgs:version1
~~~

After download and installation, the command opens a Unix terminal running in the Docker container so users can begin their analysis. From the Unix terminal running in the Docker container, users need only execute the following command to run the MutantHuntWGS pipeline:

~~~
$ MutantHuntWGS.sh \
-n FILENAME \
-g /Main/MutantHuntWGS/S_cerevisiae_Bowtie2_Index_and_FASTA/genome \
-f /Main/MutantHuntWGS/S_cerevisiae_Bowtie2_Index_and_FASTA/genome.fa \
-r single \
-s 100 \
-p /Main/MutantHuntWGS/S_cerevisiae_Bowtie2_Index_and_FASTA/ploidy_n1.txt \
-d /Main/Analysis_Directory \
-o /Main/Analysis_Directory/NAME_YOUR_OUTPUT_FOLDER
-a YES
~~~

A detailed description of the pipeline, its installation, and its use is available in the MutantHuntWGS Git repository readme (https://github.com/mae92/MutantHuntWGS/blob/master/README.md).

### Utility of the MutantHuntWGS Pipeline

MutantHuntWGS processes WGS data through a standard alignment/variant-calling pipeline and compares each experimental strain to a control strain (Figure 1, see Methods). The pipeline’s constituent tools are often used for WGS analysis (Reavey *et al*. 2015; Gopalakrishnan and Winston 2019; Gopalakrishnan *et al*. 2019). However, MutantHuntWGS ensures ease of use by assembling these tools in a Docker container and requiring only one command to run them all in sequence. This approach combines the best aspects of previously published pipelines (discussed below) while allowing inexperienced users to install the software and reproducibly apply popular methods.

MutantHuntWGS also ensures that the output data files are well organized and easy to locate. Output files include aligned reads (BAM format), alignment statistics (TXT format), pre- and post-filtering variants (VCF format), SnpEff output (HTML, VCF, and TXT formats), and SIFT output (VCF, XLS formats). The user thus has all the information needed to identify and visually inspect sequence variants, and to generate figures and tables for publication.

### MutantHuntWGS combines versatility and simplicity

Our goal in creating MutantHuntWGS was to simplify the installation and usage of robust bioinformatics tools while maintaining flexibility by allowing users to specify certain critical options. Examples of this, discussed below, include (1) enabling use with additional organisms, (2) allowing users to specify ploidy, (3) filtering by a user-specified variant-quality score, and (4) exposing all intermediate and final output files to facilitate additional filtering and quality control.

MutantHuntWGS is designed for use with *S. cerevisiae* by default but can be adapted to analyze WGS data from any organism. At present, only the necessary reference files for *S. cerevisiae* are included in the MutantHuntWGS download. Investigators who wish to analyze data from an organism other than *S. cerevisiae* need to provide, at minimum, new Bowtie2 indices, a genome FASTA file, and a ploidy file. Bowtie2 indices and genome FASTA files for many model organisms are available at https://support.illumina.com/sequencing/sequencing_software/igenome.html. A FASTA index file (genome.fasta.fai) that can be easily converted into a ploidy file is also available at this link. Unfortunately, performing the SnpEff and SIFT analysis would require slight alterations to the SnpEff and SIFT commands in the pipeline script and a copy of the SIFT library for the organism of interest. We chose not to include reference files and SIFT libraries for other organisms within the Docker container due to the large size of these files. If users encounter difficulties when analyzing non-*S. cerevisiae* WGS data, we encourage them to seek assistance by opening an issue on the MutantHuntWGS Git repository.

Experiments in yeast are often performed in a haploid background, but can also be performed in diploid or occasionally aneuploid backgrounds. The MutantHuntWGS download includes two ploidy files, one for diploids and one for haploids. The user can specify either ploidy file when running the pipeline. MutantHuntWGS will automatically provide this file to BCFtools during the variant-calling step.

Users may also set variant-quality-score cutoffs (described in detail on GitHub: https://github.com/mae92/MutantHuntWGS/blob/master/README.md) to tune the stringency of the analysis. They can also toggle the alignment step to save time when resetting the stringency. This option re-subsets variant calls with a higher or lower stringency cutoff, skipping the more time-consuming upstream steps of the pipeline. Although MutantHuntWGS does not allow users to specify additional cutoffs that filter the output per SnpEff/SIFT effect predictions and scores, users can separately apply such filters to the MutantHuntWGS output files after the fact—thus allowing for increased stringency.

### Assessing MutantHuntWGS performance using a bulk segregant analysis dataset

To assess MutantHuntWGS performance, we applied it to bulk segregant analysis data (Birkeland *et al*. 2010) with ploidy set to haploid. MutantHuntWGS identified 188 variants not present in the control strain that passed the variant-quality-score cutoff of 100. Thus only 1.95% of all variants detected in the experimental strain passed the filtering steps (Table 1). Among these was the same *PHO81* (*VAC6*) mutation found in the Birkeland *et al*. (2010) study, which results in an R701S amino acid substitution in the Pho81 protein (Birkeland *et al*. 2010). Our pipeline thus identified the same published causal variant described in the original study.

**Table 1.**
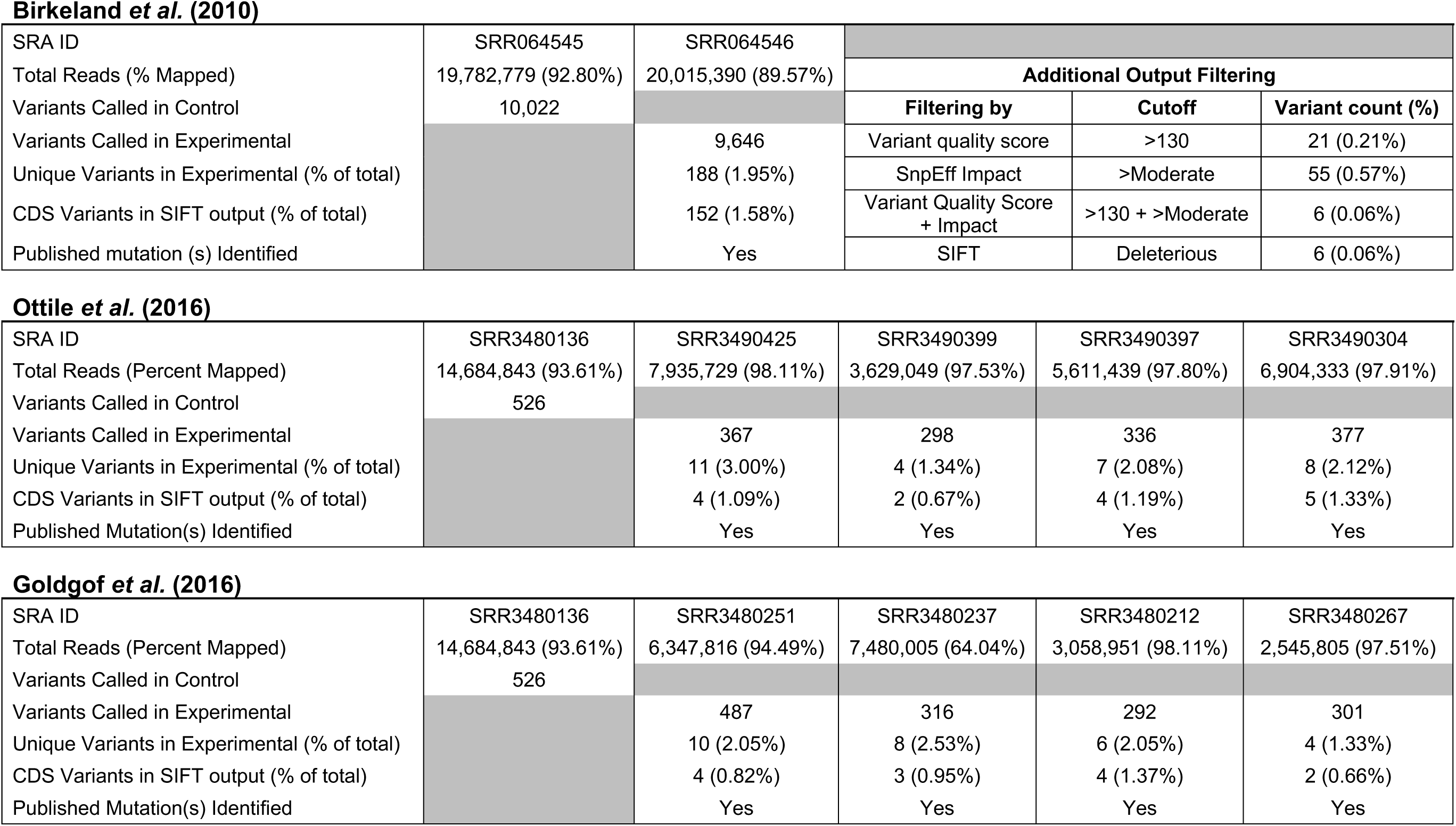
Analysis of previously published bulk-segregant and lab-evolution WGS datasets using MutantHuntWGS demonstrates the effectiveness of the pipeline.

**Birkeland *et al.* (2010)**
|  |  |  |  |  |  |
| --- | --- | --- | --- | --- | --- |
| SRA ID | SRR064545 | SRR064546 |  |  |  |
| Total Reads (% Mapped) | 19,782,779 (92.80%) | 20,015,390 (89.57%) | **Additional Output Filtering** |  |  |
| Variants Called in Control | 10,022 |  | **Filtering by** | **Cutoff** | **Variant count (%)** |
| Variants Called in Experimental |  | 9,646 | Variant quality score | >130 | 21 (0.21%) |
| Unique Variants in Experimental (% of total) |  | 188 (1.95%) | SnEff Impact | >Moderate | 55 (0.57%) |
| CDS Variants in SIFT output (% of total) |  | 152 (1.58%) | Variant Quality Score + Impact | >130 + >Moderate | 6 (0.06%) |
| Published mutation (s) Identified |  | Yes | SIFT | Deleterious | 6 (0.06%) |
| **Ottile *et al.* (2016)** |  |  |  |  |  |
| SRA ID | SRR3480136 | SRR3490425 | SRR3490399 | SRR3490397 | SRR3490304 |
| Total Reads (Percent Mapped) | 14,684,843 (93.61%) | 7,935,729 (98.11%) | 3,629,049 (97.53%) | 5,611,439 (97.80%) | 6,904,333 (97.91%) |
| Variants Called in Control | 526 |  |  |  |  |
| Variants Called in Experimental |  | 367 | 298 | 336 | 377 |
| Unique Variants in Experimental (% of total) |  | 11 (3.00%) | 4 (1.34%) | 7 (2.08%) | 8 (2.12%) |
| CDS Variants in SIFT output (% of total) |  | 4 (1.09%) | 2 (0.67%) | 4 (1.19%) | 5 (1.33%) |
| Published Mutation(s) Identified |  | Yes | Yes | Yes | Yes |
| **Goldgof *et al.* (2016)** |  |  |  |  |  |
| SRA ID | SRR3480136 | SRR3480251 | SRR3480237 | SRR3480212 | SRR3480267 |
| Total Reads (Percent Mapped) | 14,684,843 (93.61%) | 6,347,816 (94.49%) | 7,480,005 (64.04%) | 3,058,951 (98.11%) | 2,545,805 (97.51%) |
| Variants Called in Control | 526 |  |  |  |  |
| Variants Called in Experimental |  | 487 | 316 | 292 | 301 |
| Unique Variants in Experimental (% of total) |  | 10 (2.05%) | 8 (2.53%) | 6 (2.05%) | 4 (1.33%) |
| CDS Variants in SIFT output (% of total) |  | 4 (0.82%) | 3 (0.95%) | 4 (1.37%) | 2 (0.66%) |
| Published Mutation(s) Identified |  | Yes | Yes | Yes | Yes |

We were surprised by how many sequence variants (relative to the reference genome) remained after filtering. Given our variant-quality-score cutoff of 100, it is unlikely that these variants were called in error; instead, they likely reflect high sequence heterogeneity in the genetic backgrounds of the experimental and control strains. To further reduce the length of the variant list, we experimented with additional cutoffs, including (1) more stringent variant-quality-score, (2) SIFT score, and (3) SnpEff impact score cutoffs. A SIFT-score cutoff of <0.05 (deleterious) reduced the number of variants in the SIFT output from 152 to 6 (Table 1). An increased variant-quality-score stringency (> 130) reduced the number of variants to 21. A SnpEff impact-score cutoff of > Moderate reduced the number of variants to 55. Finally, a variant quality-score cutoff of > 130 and a SnpEff score of > Moderate, used together, reduced the number of variants to only 6. All post-hoc tests retained the causal variant. These tests demonstrate how users might similarly narrow their lists of potential candidates.

### Assessing MutantHuntWGS performance using lab evolution datasets

To test MutantHuntWGS performance on strains that did not undergo bulk-segregant analysis, we analyzed nine datasets from lab evolution experiments (Goldgof *et al*. 2016; Ottilie *et al*. 2017), again setting ploidy to haploid and using a variant-quality-score cutoff of 100. In each of these studies, yeast cells were allowed to evolve resistance to a drug and WGS was used to identify mutations (Goldgof *et al*. 2016; Ottilie *et al*. 2017).

Shortlists of only 4 to 11 (1.33% to 3.00%) of the variants originally detected in the experimental strain(s) were obtained for each dataset. Out of these variants, only 2 to 5 (0.66% - 1.37% of called variants in the experimental strain) were present in the SIFT output for each dataset. In each case, the list of variants generated by MutantHuntWGS included the mutation identified in the published study (SRR3480237: Pma1 N291K, Yrr1 T623K; SRR3480212: Pma1 P339T, Yrr1 L611F; SRR3480251: Pma1 L290S, Yrr1 T623K; SRR3480267: Pma1 G294S; SRR3490304: Erg11 V154G; SRR3490397: Erg11 T318N; SRR3490399: Erg25 D234E; SRR3490425: Erg25 H156N). These test cases confirm that MutantHuntWGS can identify yeast-sequence variants from WGS sequencing samples and accurately filter out background mutations.

### Existing WGS analysis pipelines

Other platforms exist that perform similar analyses. Each possesses a subset of the features enabled by MutantHuntWGS and has notable strengths. MutantHuntWGS is unique in its ability to combine the best attributes of these published tools while including additional functionality and providing output data in standard formats, such as BAM and VCF.

One user-friendly program, Mudi (Iida *et al*. 2014), uses BWA (Jo and Koh 2015), SAMtools (Li *et al*. 2009), and ANNOVAR (Wang *et al*. 2010) for sequence alignment, identification, and annotation of sequence variants, respectively. Like MutantHuntWGS, Mudi performs numerous filtering steps before returning a list of putative causal variants. MutantHuntWGS predicts variant effects and maps variants to annotated *S. cerevisiae* genes using SnpEff and SIFT instead of ANNOVAR, and also offers access to all intermediate data files.

Another program, VAMP, consists of a series of Perl scripts that build and query an SQL database made from user-provided short-read sequencing data. VAMP identifies sequence variants, including large insertions and deletions. It also has built-in functionality that allows for manual inspection of the data (Birkeland *et al*. 2010). One advantage of MutantHuntWGS over VAMP is that it adheres to common data formats.

A recent article describing WGS in yeast samples includes a bioinformatics pipeline, referred to as wgs-pipeline (Gopalakrishnan and Winston 2019). It is built in a Snakemake framework (Köster and Rahmann 2012) that runs in a Conda environment (https://docs.conda.io/en/latest/), similar to the container-based analysis environment we used for MutantHuntWGS. This pipeline uses Bowtie2 (Langmead and Salzberg 2012), SAMtools (Li *et al*. 2009), Picard (Toolkit 2016), and GATK (McKenna *et al*. 2010) to align, process, and compare datasets. Compared to wgs-pipeline, MutantHuntWGS, which runs both SnpEff and SIFT on the candidate variants, provides a more comprehensive analysis of the predicted effects of the variants.

The Galaxy platform (Giardine *et al*. 2005; Blankenberg *et al*. 2010) provides a user-friendly, online interface for building bioinformatics pipelines. Galaxy also offers access to intermediate files. However, analysis with this platform requires the user to select the tools and parameters to incorporate, so some knowledge of the tools themselves is essential. Implementation is straightforward after those decisions are made, and the user need not have any understanding of Unix/Linux. The advantage of MutantHuntWGS over the Galaxy platform and pipelines such as CloudMap (Minevich *et al*. 2012) is that the user does not need to make decisions about the data analysis workflow.

In summary, the MutantHuntWGS pipeline is among the most user-friendly of these programs. It combines the most useful features of the existing WGS analysis programs while also enabling the user to account for ploidy. Containerization streamlines the installation of MutantHuntWGS and enhances its reproducibility. Thus, MutantHuntWGS offers ease of use, functionality, and data-transparency, setting it apart from other WGS pipelines.

### Conclusions

Processing data generated from next-generation sequencing platforms requires significant expertise, and so is inaccessible to many investigators. We have developed a highly effective differential variant-calling pipeline capable of identifying causal variants from WGS data. We demonstrate the utility of MutantHuntWGS by analyzing previously published datasets. In all cases, our pipeline successfully identified the causal variant. We offer this highly reproducible and easy-to-implement bioinformatics pipeline to the *Saccharomyces cerevisiae* research community (available at https://github.com/mae92/MutantHuntWGS).

## ACKNOWLEDGMENTS

We would like to thank Margaret Shirra, Sarah Tripplehorn, Alex Francette, and Brendan McShane for careful review of the manuscript. This research was supported by a National Institutes of Health (NIH) grant R01GM052593 to KMA, a predoctoral fellowship from the NIH (F31GM129917) awarded to MAE, and University of Pittsburgh Central Research Development Funds (CRDF, 2017-2018) to JDD.

## SUPPLEMENTAL METHODS

### Pipeline Overview

The MutantHuntWGS pipeline integrates a series of open-source bioinformatics tools and Unix commands that accept raw sequencing reads (compressed FASTQ format or .fastq.gz) and a text file containing ploidy information as input, and produces a list of sequence variants as output. The user must provide input data from at least two strains: a control strain and one or more experimental strains. The pipeline uses (1) Bowtie2 to align the reads in each input sample to the reference genome (Langmead and Salzberg 2012), (2) SAMtools to process the data and calculate genotype likelihoods (Li *et al*. 2009), (3) BCFtools to call variants (Li *et al*. 2009), (4) VCFtools (Danecek *et al*. 2011) and custom shell commands to compare variants found in experimental and control strains, and (5) SnpEff (Cingolani *et al*. 2012) and SIFT (Vaser *et al*. 2016) to assess where variants are found in relation to annotated genes and the potential impact on the expression and function of the affected gene products (see Figure 1 in main paper).

### Sequence Alignment

MutantHuntWGS uses Bowtie2 version 2.2.9 (Langmead and Salzberg 2012) to first align the raw reads present in the input FASTQ files to the *S. cerevisiae* genome (S288C version = R64-2-1) (Cherry *et al*. 2012; Engel *et al*. 2014). As the default, we set Bowtie2 to search for a maximum of two distinct alignments per read `(-k 2` option), which reduces alignment time and multiple mapping. The pipeline only retains sequencing reads that align (`--no-unal` option) in the SAM (Sequence Alignment/Map Format) output to help reduce file size. MutantHuntWGS uses SAMtools version 1.3.1 to convert the aligned-read output from Bowtie2 (SAM format) into the BAM (Binary Alignment/Map) format (view -bS options) (Li *et al*. 2009). SAMtools then sorts (`sort` option) and indexes (`index` option) the BAM file to prepare the data for variant calling. Users can view the sorted and indexed BAM files in a genome browser such as IGV (Integrative Genome Viewer) to examine the aligned reads (Thorvaldsdottir *et al*. 2013).

### Variant Calling

Based on the aligned reads, SAMtools outputs genotype likelihoods as BCF (Binary Call Format) files (`mpileup -g -f` options) using the BAM file as input (Li *et al*. 2009). BCFtools version 1.3.1 then uses the genotype likelihoods recorded in the BCF file to call single nucleotide polymorphisms (SNPs), as well as insertions and deletions (INDELs) (`-c -v --samples-file –ploidy-file` options) (Li 2011). This variant information is saved in the Variant Call Format (VCF), the format used by the 1000 Genomes Project (Danecek *et al*. 2011). IGV can again be used to view the VCF files that are output from MutantHuntWGS (Thorvaldsdottir *et al*. 2013). At the variant calling step, BCFtools also considers a user-specified input ploidy file to account for genome copy number.

### Identifying Candidate Variants

VCFtools version 0.1.14 compares VCF files (`--diff-site` option) from the control and experimental samples (Danecek *et al*. 2011). To retain the variants that are found only in the experimental dataset, MutantHuntWGS uses the Unix `awk` command (Aho *et al*. 1979) to remove variants from the VCFtools output that have VCF scores lower than a user-defined variant-quality-score cutoff. For each experimental dataset, it then uses the Unix `grep`, `head`, and `cat` commands to construct new VCF files that contain only the variants specific to the experimental strain. These VCF files can also be viewed in IGV.

### Variant Effect Prediction

SnpEff version 4.3p (Cingolani *et al*. 2012) and SIFT4G (i.e., SIFT) (Vaser *et al*. 2016) are useful programs for (1) determining whether sequence variants are located in or near an annotated coding region and (2) predicting the effect the variant might have on gene expression or function of the protein product. SnpEff determines the locations of sequence variants relative to protein-coding genes and the severity of each variant based on how likely it is to disrupt gene expression or function (Cingolani *et al*. 2012).

SnpEff also annotates variants in 5’ and 3’ UTRs as well as promoter regions. This information is vital if the causal mutation disrupts a ncRNA or DNA element rather than altering a protein-coding sequence. SIFT uses the EF4.74 library for *S. cerevisiae* to score variants found in protein-coding genes in order to predict the impact of the resulting amino acid changes (Vaser *et al*. 2016). MutantHuntWGS saves all SnpEff and SIFT output files so the user can further filter the results to reduce the number of candidate sequence variants identified.

### Analysis of previously published data

To demonstrate utility, we used MutantHuntWGS to analyze published datasets from paired-end sequencing experiments with DNA prepared from bulk segregants or lab-evolved strains (Birkeland *et al*. 2010; Goldgof *et al*. 2016; Ottilie *et al*. 2017). These data were downloaded from the sequence read archive (SRA) database (https://www.ncbi.nlm.nih.gov/sra; project accessions: SRP003355, SRP074482, SRP074623) and decompressed using the SRA toolkit (https://github.com/ncbi/sra-tools/wiki). MutantHuntWGS was run from within the Docker container, and each published mutant (experimental) file was compared to its respective published control. When processing data from bulk segregant analysis, we reduced the number of candidate variants by additionally using more stringent cutoffs: variant quality score > 130, SnpEff impact score > Moderate, and SIFT score < 0.05 (deleterious).

